# Genome-wide admixture mapping of eGFR and CKD identify European and African ancestry-of-origin loci in U.S. Hispanics/Latinos

**DOI:** 10.1101/2021.05.06.442996

**Authors:** Andrea R.V.R. Horimoto, Jianwen Cai, James P. Lash, Martha L. Daviglus, Nora Franceschini, Timothy A. Thornton

## Abstract

**Background:** Admixture mapping is a powerful approach for gene mapping of complex traits that leverages the diverse genetic ancestry in populations with recent admixture such as U.S. Hispanics/Latinos (HL), who have increased risk of chronic kidney disease (CKD).

**Methods:** Genome-wide admixture mapping was performed for CKD and estimated glomerular filtration rate (eGFR) in a sample of 12,601 participants from the Hispanic Community Health Study/Study of Latinos, with validation in a sample of 8191 African Americans from the Women’s Health Initiative (WHI).

**Results:** Three novel ancestry-of-origin loci were identified on chromosomes 2, 14 and 15 for CKD and eGFR. The chromosome 2 locus (2p16.3) consisted of two European ancestry regions encompassing the *FSHR* and *NRXN1* genes, with European ancestry at this locus associated with increased risk for CKD. The chromosome 14 locus (14q32.2) located within the *DLK1-DIO3* imprinted domain was driven by European ancestry, and was associated with lower eGFR. The chromosome 15 locus (15q13.3-14) included intronic variants of *RYR3* and was within an African-specific genomic region that was associated with higher eGFR. These findings were compared to the conventional genome-wide association study that failed to identify significant associations in these regions. We validated the chromosome 14 and 15 loci for eGFR in the WHI African Americans.

**Conclusions:** This study provides evidence of shared ancestry-specific genomic regions influencing eGFR in HL and African Americans, and illustrates the potential for leveraging genetic ancestry in recently admixed populations for novel discovery of kidney trait loci.

## Introduction

Chronic kidney disease (CKD) affects one in seven U.S. adults, with a disproportionate burden on African Americans (AA) and Hispanics/Latinos (HL).^1,2^ Ancestry-specific genetic contributions to CKD risk have been identified in AA and HL, such as the African-derived *APOL1* G1 and G2 genotypes.^3,4^ Few genome-wide association studies (GWAS) or multi-ethnic GWAS meta-analysis of CKD and estimated glomerular filtration rate (eGFR) have included HL populations.^5–11^ Since genetic risk may not be shared across populations^12^, leveraging the genetic diversity in recently admixed populations offers an opportunity to identify novel loci that have not been detected in genetic studies of homogeneous ancestry populations.

Admixture mapping is a powerful approach for genetic mapping of complex diseases in multi-ethnic populations. Recent admixture between continental ancestral populations, such as in HL, creates long-range blocks of correlation between genetic variants in chromosome segments (linkage disequilibrium, LD) for variants that have large allele frequency differences in the founding populations. Therefore, the genome of an admixed individual is a mosaic of segments of different ancestral origin (**Figure 1A**). The rationale for admixture mapping is that causal variants have a higher frequency in segments inherited from ancestral populations with higher disease prevalence. Admixture mapping leverages the genetic ancestry to identify associations between these ancestry-specific chromosomal segments (local ancestry) at each locus and traits, yielding new and complementary findings compared to GWAS (association mapping) of complex diseases. Admixture mapping led to the discovery of the *APOL1* genomic region for focal segmental glomerulosclerosis and hypertension-attributed end-stage kidney disease in AA. Subsequent fine-mapping within the chromosome 22 region identified the G1 and G2 risk genotypes.^3^

**Figure 1.**
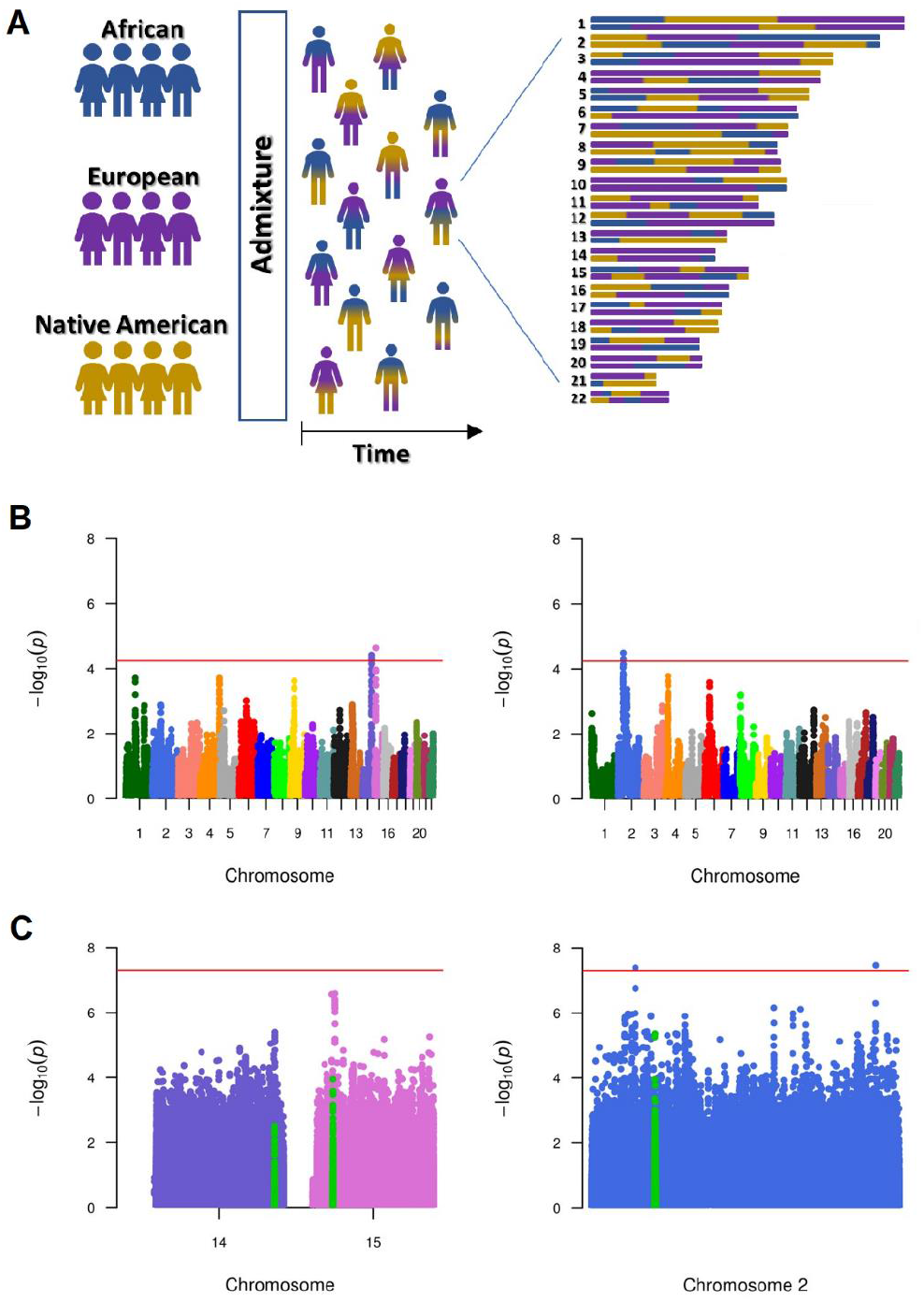
Recent admixture representation of Hispanic/Latinos populations (**A**), admixture mapping Manhattan plots for eGFR (at left) and CKD (at right) in HCHS/SOL (**B**), and GWAS Manhattan plots for eGFR (at left) and CKD (at right) in HCHS/SOL (**C**). Only GWAS results for the chromosomes of interest 2, 14 and 15 are represented. SNPs in the regions associated in the admixture mapping are highlighted in green. Note two significant signals on chromosome 2 unrelated to the admixture mapping findings in the GWAS for CKD driven by low frequency variants (rs146424372 and rs116137931).

HL populations are admixed with European, African, and Native American ancestries. Using an admixture mapping approach that allows for the testing of multiple ancestry effects simultaneously, our recent studies identified a novel American Indian locus at chromosome 2 associated with urine albumin excretion.^13^ In addition to identifying trait-associated loci with causal variants that have large allele frequency differences across ancestries, our approach also allows for the identification of the ancestries that are driving these associations. We also recently extended our admixture mapping approach for analyses of dichotomous outcomes such as CKD.^14^ This study aims to identify and fine-map kidney loci for eGFR and CKD using admixture mapping in HL participants of the Hispanic Community Health Study/Study of Latinos (HCHS/SOL).

## Methods

### Study samples and phenotype

HCHS/SOL is a community-based cohort study that recruited 16,415 participants from four U.S. field centers (Bronx, Chicago, Miami, and San Diego) representative of the largest U.S. HL groups: Central Americans, South Americans, Cubans, Mexicans, Puerto Ricans, and Dominicans. The study has extensive familial relatedness, as previously discussed.^15^ Details about the complex sampling design and cohort selection for HCHS/SOL are described elsewhere.^16^ Physical, sociodemographic and lifestyle evaluations, and fasting blood and spot urine samples were collected in the baseline (2008-2011). The HCHS/SOL study was approved by the institutional review board at each participating institution and collected a written informed consent from all participants.

Participants were genotyped for more than 2.5 million single nucleotide polymorphisms (SNPs) using a customized Illumina array, and imputation was performed using the 1000 Genomes Project phase I reference panel. The quality control procedures and details about genotyping, imputation and local ancestry calls were previously described.^15,17,18^ eGFR, estimated using the Chronic Kidney Disease Epidemiology Collaboration equation^19^ based on serum creatinine measured using a creatinase enzymatic method, was inverse normal transformed for validity of the normality assumptions for the statistical procedures. CKD was defined by an untransformed eGFR < 60 ml/min/1.73 m^2^. We performed analyses for eGFR and CKD in a subset of 12,601 subjects with available genotype and phenotype data.

An independent sample of 3050 HL and 8191 AA women from the Women’s Health Initiative^20^ SNP Health Association Resource (WHI SHARe) were used to validate our findings. Local ancestry calls for both WHI HLs and AAs were inferred previously.^21^

## Statistical Analysis

### Multiple random and fixed effects

The HCHS/SOL study was developed under a complex sampling design. To account for the correlation structure of the data, we included three random effects: the pairwise kinship coefficients, household, and census block group (the geographic cluster of the households), which represent the polygenic effects due to the familial structure and shared environment effects. The statistical model was additionally fitted for the first five principal components that have been shown to account for population structure in our data, recruitment center, genetic-analysis group, age, and sex as fixed effects. The genetic-analysis groups correspond to the same self-identified background groups (Central American, South American, Cuban, Mexican, Puerto Rican or Dominicans), where samples were relocated to compound more genetically homogenous clusters. Details on the estimation of genetic analysis groups, principal components and kinship coefficients were previously reported.^15^ For linear mixed models (LMM), we also estimated separate residual variance components by genetic-analysis group to avoid violations of the homoscedasticity assumptions.

### Admixture mapping

The admixture mapping was performed using LMM and logistic mixed model (GMMAT^14^) for eGFR and CKD outcomes, respectively, in which all European, African and Native American ancestries were tested simultaneously. Statistical details about LMM and GMMAT are provided in **Supplemental Appendix 1**. Admixture mapping with LMM and GMMAT are implemented in the GENESIS R package^22^. We first fit the models under the null hypothesis of no genetic effect, including the random and fixed effects described above. We then used the null models to evaluate the association between the local ancestry at each locus and the outcomes using a score test. Here, locus is defined as a long-range block of admixture-linkage disequilibrium created by the recent admixture in HL and local ancestry correspond to the locus-specific ancestry allelic dosages (0, 1 or 2 copies of European, African, or Native American alleles) estimated from the genotype data. As every SNP in a locus has identical association signal and effect size, only one SNP per locus, here named as lead SNP, was included in the analyses totalizing 15,500 loci tested. Secondary admixture mapping analyses were performed by testing each ancestry against the others to identify which ancestry accounts for the association at each locus. Based on previous simulation analysis, a nominal *P* of 5.7 x 10^−5^ controls the type I error at level of 0.05^23^, and accounts for the number of independent tests performed under populational structure. Given the long-range LD structure in recent admixed populations, a smaller number of tests should be conducted and, consequently, significance thresholds applied for admixture mapping are not as stringent as in GWAS. The effect sizes for the significant ancestry blocks were estimated based on the allelic dosage of the ancestry driving the signal. To provide evidence for the locus accounting for the observed local ancestry association, we conducted conditional admixture mapping including the lead SNPs representing each associated locus as covariate in models.

### Association Mapping (GWAS)

We performed a GWAS on the HCHS/SOL imputed genotype data using the GENESIS R package^22^. We first filtered imputed SNPs based on the information metric provided by IMPUTE2^24^ (info > 0.8), the ratio of observed variance of imputed dosages to the expected binomial variance (oevar > 0.8), and the effective minor allele count (2p(1-p)*N*oevar, where p is the minor allele frequency and N is the sample size) > 30 for eGFR and >50 for CKD. Using the null models fit for the admixture mapping, we tested the association between each SNP, using the dosage of reference allele, and the outcomes using a score test.

### Admixture mapping validation

We performed validation studies for the HCHS/SOL admixture mapping findings in 3,050 HL and 8,191 AA independent samples from the Women’s Health Initiative (WHI) SNP Health Association Resource (WHI SHARe)^20^, using the local ancestry calls previously estimated.^21^ Admixture mapping for eGFR and CKD outcomes in HL and AA samples were conducted in the regions of interest using mixed models adjusted by age and top four PCs. Admixture mapping and coefficients estimation for the associated loci followed the methods described above.

## Data sharing

The Hispanic Community Health Study / Study of Latinos (HCHS/SOL) data supporting the findings of this study is openly available in the dbGap repository at https://www.ncbi.nlm.nih.gov/gap/, under the accession numbers phs000880.v1.p1 and phs000810.v1.p1.

## Results

Table 1 displays the characteristics of HCHS/SOL participants including comorbidities. The mean age was 46.1 (range 18 to 76 years), and 59% were females. The overall CKD prevalence based on an eGFR <60 ml/min/1.73 m^2^ was 3.4%.

**Table 1.**
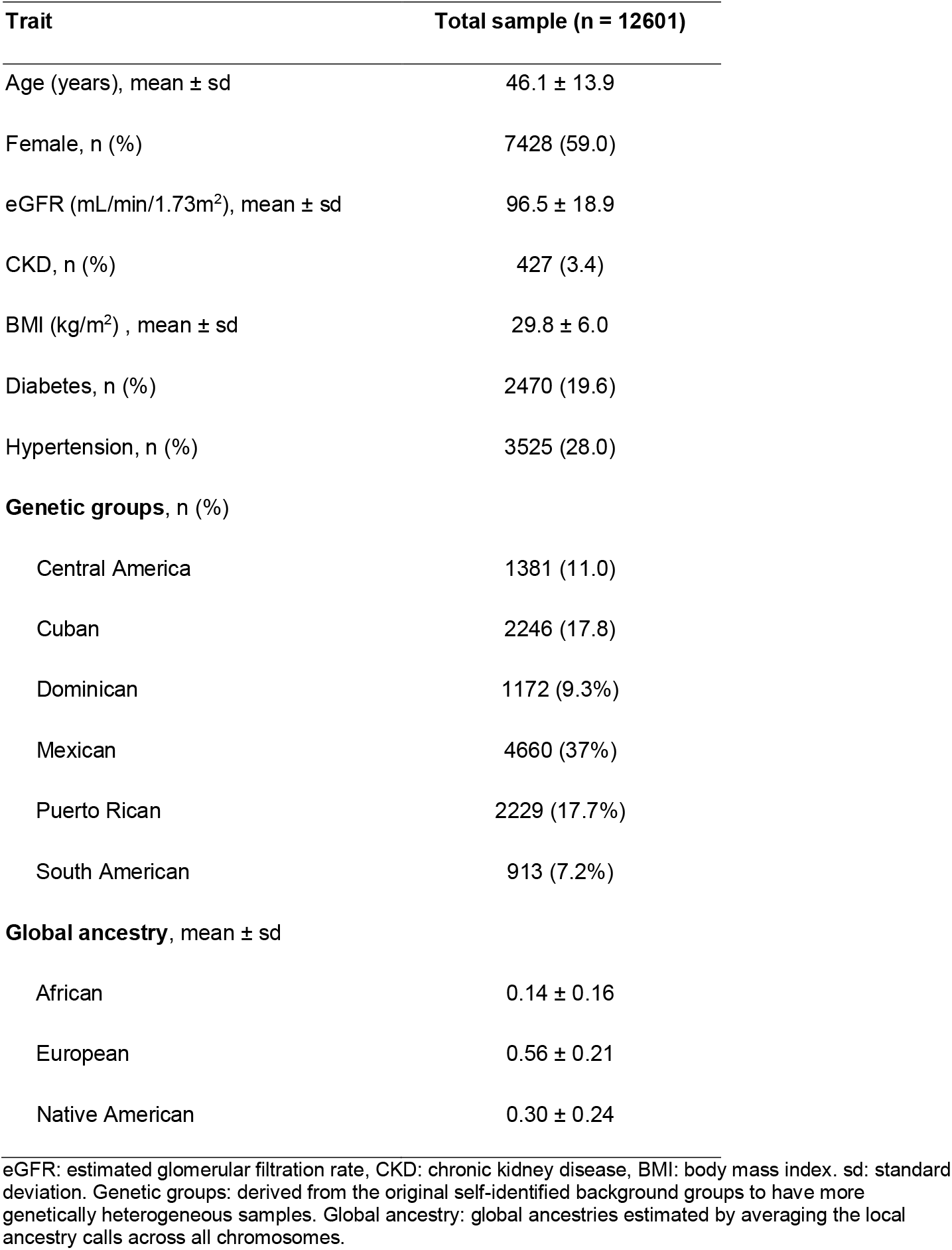
Descriptive characteristics of HCHS/SOL samples.

| **Trait** | **Total sample (n = 12601)** |
| --- | --- |
| Age (years), mean ± sd | 46.1 ± 13.9 |
| Female, n (%) | 7428 (59.0) |
| eGFR (mL/min/1.73m<sup>2</sup>), mean ± sd | 96.5 ± 18.9 |
| CKD, n (%) | 427 (3.4) |
| BMI (kg/m<sup>2</sup>) , mean ± sd | 29.8 ± 6.0 |
| Diabetes, n (%) | 2470 (19.6) |
| Hypertension, n (%) | 3525 (28.0) |
| **Genetic groups, n (%)** |  |
| Central America | 1381 (11.0) |
| Cuban | 2246 (17.8) |
| Dominican | 1172 (9.3%) |
| Mexican | 4660 (37%) |
| Puerto Rican | 2229 (17.7%) |
| South American | 913 (7.2%) |
| **Global ancestry, mean ± sd** |  |
| African | 0.14 ± 0.16 |
| European | 0.56 ± 0.21 |
| Native American | 0.30 ± 0.24 |

eGFR: estimated glomerular filtration rate, CKD: chronic kidney disease, BMI: body mass index. sd: standard deviation. Genetic groups: derived from the original self-identified background groups to have more genetically heterogeneous samples. Global ancestry: global ancestries estimated by averaging the local ancestry calls across all chromosomes.

### Admixture mapping findings for eGFR and CKD

The admixture mapping of eGFR identified two genome-wide significant loci on chromosomes 14 and 15, while the admixture mapping of CKD identified two loci on chromosome 2 (**Figure 1B**). When testing each ancestry contribution to admixture mapping findings, the association on chromosome 14 was driven by European ancestry, while African ancestry was driving the association on chromosome 15 (**Figure 2A**). For the CKD loci, the chromosome 2 associations were driven by European ancestry (**Figure 2B**). Therefore, European and African ancestry chromosome segments contributed to our new findings (**Table 2**).

**Figure 2.**
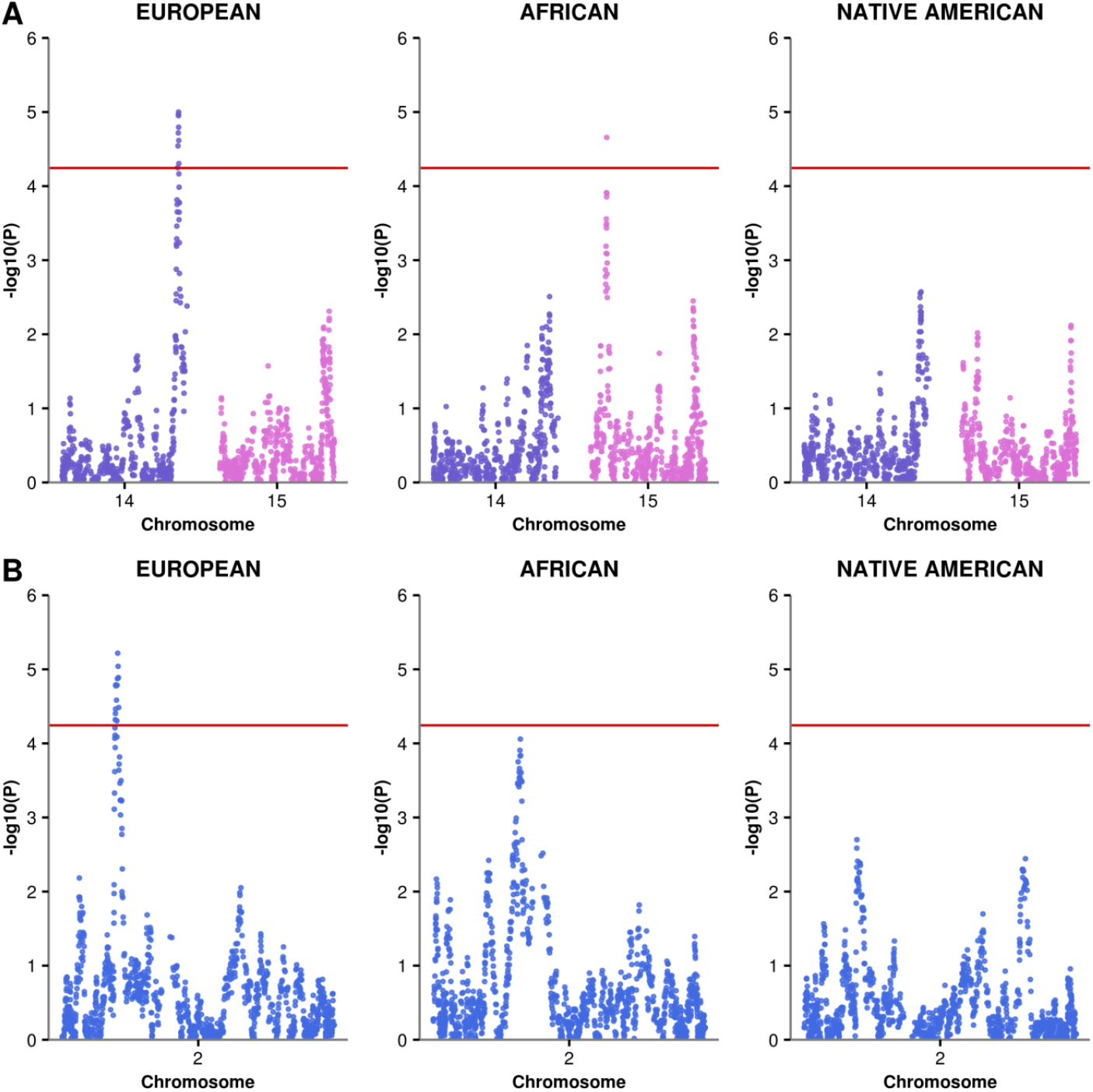
Single ancestry admixture mapping analyses for eGFR (**A**) and CKD (**B**), in which European, African, and Native American ancestries were tested separately. Note that the eGFR chromosome 14 signal is driven by European ancestry and the chromosome 15 signal is driven by African ancestry. For CKD, the chromosome 2 signal is driven by European ancestry.

**Table 2.** Admixture mapping results for eGFR and CKD.

| HCHS/SOL admixture mapping results |  |  |  |  |  |  | WHI African Americans admixture mapping validation |  |  |  |  |
| --- | --- | --- | --- | --- | --- | --- | --- | --- | --- | --- | --- |
| Trait | Chr | #SNPs | Physical position<br>(GRCh37/hg19) | AM joint<br>*P* | Effect<br>(95% CI) | Ancestry<br>background |  | #SNPs | Physical position<br>(GRCh37/hg19) | AM joint<br>*P* | Effect<br>(95% CI) |
|  |  |  |  |  |  | Ancestry | *P* |  |  |  |  |
| eGFR | 14 | 4 | 101329926 - 101350298 | 4.0 x 10<sup>-5</sup> | -0.05<br>(-0.07; -0.03) | EUR | 1.0 x 10<sup>-5</sup> | 2 | 101349017 -101350298 | 0.02 | -0.04<br>(-0.08; -0.01) |
|  | 15 | 37 | 33801207 - 33884488 | 2.4 x 10<sup>-5</sup> | 0.07<br>(0.04; 0.11) | AFR | 2.2 x 10<sup>-5</sup> | 5 | 33801207-33812915 | 0.03 | -0.04<br>(-0.08; -0.003) |
|  |  |  |  |  |  |  |  | 41 | 33821715-33884488 | 0.04 | -0.04<br>(-0.08; -0.002) |
| CKD | 2 | 66 | 49571948 - 49894154 | 3.3 x 10<sup>-5</sup> | 1.46<br>(1.23; 1.73) | EUR | 6.1 x 10<sup>-6</sup> |  | No validation |  |  |
|  |  | 37 | 49909155 - 50197075 | 5.1 x 10<sup>-5</sup> | 1.45<br>(1.23; 1.71) | EUR | 9.1 x 10<sup>-6</sup> |  | No validation |  |  |

Chr: chromosome; #SNPs: number of SNPs within each locus; AM joint *P*: *P* for the admixture mapping joint test, in which all ancestries were tested simultaneously. The significance threshold for admixture mapping is a $P < 5.7 \times 10^{-5}$ ; Effect (95%CI): coefficient (95% confidence interval) for eGFR and odds ratio (95% confidence interval) for CKD. Ancestry background: ancestry driving the signal, according to the single ancestry admixture mapping analysis, and the respective *P* of the statistical test.

The eGFR association on chromosome 14 was captured by four SNPs. The beta coefficient for local ancestry was −0.05 (standard error [SE] = 0.01, *P* = 9.8 x 10^−6^) indicating a decreasing on eGFR for each additional copy of the European allele at this locus. The eGFR association on chromosome 15 was observed in a locus with 37 SNPs, all of them within the *RYR3* gene, where African ancestry conferred a protective effect against low eGFR (beta coefficient = 0.07, SE = 0.02, *P* = 2.1 x 10^−5^).

The CKD association on chromosome 2 was driven by two loci with 66 and 37 SNPs, respectively. These regions include intergenic variants nearby the *FSHR* and *NRXN1* genes and intronic variants of *NRXN1*. The odds ratio for CKD was similar across the two loci (1.46, 95% CI = 1.23 −1.73; 1.45, 95% CI = 1.23 −1.71 for the first and second locus, respectively) which increased risk of CKD for each additional copy of the European allele (**Supplementary Table 1)**. Overall, our studies have mapped three novel loci associated with eGFR or CKD in HL, which were driven by ancestry-specific background.

### Conditional admixture mapping analyses

To provide supporting evidence for identified loci accounting for observed admixture mapping associations, we performed conditional admixture mapping analyses including the lead SNP of each locus as covariate. Conditional analyses on one or more lead SNPs completely accounted for the associations on chromosomes 2, 14 and 15 (**Supplementary Figure 1)**. Note that the two regions on chromosome 2 are only 15Kb apart and show similar effect, and they explained the admixture association whether tested jointly (**Supplemental Figure 1D**) or separately (**Supplemental Figures 1E and 1F**). These findings support the presence of a single signal at these loci.

### Association mapping results (GWAS)

To compare gene discovery from admixture mapping to discovery using genome-wide association, we performed a GWAS for eGFR and CKD, fitting similar statistical models as applied to admixture mapping to avoid bias due to adjustments. The GWAS did not identify significantly associated loci at the chromosomes 2, 14 and 15 regions (**Figures 1C**). In the GWAS for eGFR, no SNP reached the genome-wide significant threshold (*P* < 5.0 x 10^−8^) on chromosomes 14 and 15. For CKD, the GWAS identified two low frequency variants (minor allele frequency = 0.3%) statistically significant on chromosome 2 (rs146424372, 25kb downstream of *SCG2, P* = 3.5 x 10^−8^; rs116137931, intronic variant of *LINC01317, P* = 4.1 x 10^−8^) in distinct regions of the admixture mapping signals. These findings support gains in gene discovery when using admixture mapping that accounts for local ancestry compared to GWAS in HL.

### Validation of the admixture mapping findings

To validate loci identified in the HCHS/SOL admixture mapping, we performed admixture mapping on the chromosomes 2, 14 and 15 regions of interest in independent HL samples from the WHI (n = 3050). We also conducted the validation on the WHI AAs (n = 8191) since the secondary admixture mapping analyses identified regions of European and African backgrounds. Both eGFR admixture mapping findings on chromosomes 14 and 15 were validated in WHI AAs (**Table 2**) but not in WHI HLs. On chromosome 14, we identified a locus with two SNPs associated with eGFR (rs6575805 and rs3825569; *P* = 0.02), in which the European background showed a similar magnitude and direction of the effect (beta coefficient = −0.04, SE = 0.02, *P* = 2.2 x 10^−2^) of that observed on HCHS/SOL. On chromosome 15, a significant association was captured by two loci including, respectively, five and 41 SNPs (*P* = 0.03 and 0.04). However, the African background in this region conferred an opposite direction of the effect observed in HCHS/SOL. The effect sizes estimated for both loci were similar (beta coefficient = −0.04, SE = 0.02, *P* = 3.3 x 10^−2^ and *P* = 4.1 x 10^−2^, respectively) suggesting that the transformed eGFR outcome decreases 0.04 for each additional copy of the African allele in this region. We did not identify a significant association in the chromosome 2 admixture mapping region of interest for CKD. In summary, we validated two loci associated with eGFR but not the chromosome 2 region associated with CKD.

## Discussion

This study is a gene discovery effort to map genetic loci for eGFR and CKD using chromosome ancestral background in HL populations. We compared our approach for gene discovery to the traditional GWAS. We demonstrated that incorporating genetic ancestry in the analysis can help to identify novel genomic regions influencing eGFR and CKD in recently admixed HL populations. The main findings of this study were the discovery of three novel loci on chromosomes 2, 14, 15 for CKD and eGFR, respectively, in which the associations were largely driven by European and African ancestries. These loci were not identified in our sample using GWAS and have not been found in large GWAS meta-analyses using ancestral homogeneous populations, despite harboring genes that been implicated in kidney diseases, as discussed below.

The European region on chromosome 2 associated with CKD comprises two loci spanning 322.2kb and 287.9kb, respectively. Variants in these loci are mostly intergenic near the *FSHR* and *NRXN1* genes (2p16.3), with some intronic to *NRXN1*, including a variant in a regulatory region (**Supplementary Table 1**). *FSHR* encodes the receptor for follicle stimulating hormone (FSH). High FSH levels were associated with the increased risk of CKD in Chinese women. In experimental models, FSH caused kidney dysfunction and tubulointerstitial fibrosis, and worsening kidney injury through macrophage recruitment and activation.^25^ The *NRXN1* gene encodes a single-pass type I membrane protein, which is involved in the central nervous system. Variants in this gene have been implicated in blood pressure, body mass index (BMI) and a range of cognitive traits.^11^

The European locus on chromosome 14 associated with eGFR includes four SNPs (**Supplementary Table 1**) spanning a 20.4 kb region within the *DLK1-DIO3* imprinted domain (14q32.2), which contains paternally expressed genes *(RTL1, DLK1, DIO3)* and maternally expressed long and small non-coding RNAs (including MEG3, *antisense RTL1*, miRNAs and pseudogenes). The *DLK1-DIO3* domain has been implicated in placental and embryonic tissues development and metabolism.^26^ A knockout mice of *Peg11/Rtl1* gene showed abnormalities of fetal capillaries leading to lethality and fetal growth retardation.^27^ All SNPs upstream/downstream of miRNAs are directly implicated on urological cancers and kidney fibrosis.^28^ MIR433 is a pro-fibrotic miRNA, and it has been considered as a potential therapeutic target for kidney fibrotic disease.^29^

The 83.2 kb African locus on chromosome 15 harbors 37 variants (35 intronic, one missense [rs2077268] and one missense/synonymous [rs674155]) in the *RYR3* gene (15q13.3-14), of which seven have regulatory features. The rs2077268 has shown a suggestive pharmacogenetic effect on heart failure among AA and HL.^30^ *RYR3* is involved in intracellular calcium ion release channels and it is expressed in skeletal muscles, brain, and kidney.^31–33^ The *RYR3* gene has been associated with diabetic kidney disease (rs2596230) in both a GWAS on AA samples and a transethnic GWAS meta-analysis, though findings did not reach genome-wide thresholds.^34^ *RYR3* has also been associated with plasma adiponectin levels and inflammation, and elevated adiponectin has been implicated with CKD occurrence and progression.^35,36^

Of the loci identified in the admixture mapping, we provided evidence for validation of the two loci for eGFR but we did not validate the chromosome 2 loci associated with CKD. The chromosome 14 locus had a consistent direction of effect for European ancestry region when comparing HCHS/SOL results and WHI. The African locus on chromosome 15 had the opposite direction of effect among HL and AA. However, the admixture mapping in WHI AA identified the same chromosome regions for eGFR supporting validation of associations even if the direction of effects was discordant. Although HL populations share European, African, and Native American background, they are highly heterogeneous in terms of proportions of ancestral populations (global ancestry proportions) and ancestry at the level of chromosomal regions (local ancestry). Admixture mapping identifies loci that be composed of multiple variants, and the locus boundaries will vary within admixed groups such as HL and AA. This may explain the negative findings in the WHI HL, as discussed below. In addition, the approach is not suitable to homogenous and not admixed populations such as individuals of European ancestry, given they lack diversity in ancestry-specific chromosome segments.

Several factors may have contributed for the lack of validation of findings in the HL sample for eGFR and in HL/AA samples for CKD, including heterogeneity in the admixture proportion and composition across studies. WHI HLs have a different ancestry composition of that observed in HCHS/SOL (**Supplementary Figure 2**). Almost half of the subjects are Mexicans (vs 37% in HCHS/SOL) and other ancestry backgrounds than Puerto Rican, Mexican and Cuban are not distinguished and likely small. WHI HL sample is also fourfold smaller than HCHS/SOL. Therefore, low statistical power, differences in genetic background, or a combination of both can explain the lack of validation of some of the regions in WHI HLs. For CKD validation, WHI AA sample was large and had higher proportion of prevalent CKD compared to the discovery sample (6% vs 3.4%). However, the frequency of

European alleles at the chromosome 2 loci was lower in WHI AA (20% for CKD cases and 24% for non-CKD in WHI AA vs 65% for CKD cases and 54% for non-CKD in HCHS/SOL). This suggests that the number of European alleles may not have captured the association at this region.

We compared our results to a GWAS and attempted to fine-map our regions using GWAS data. The GWAS did not identified genome-wide significant associations within the admixture mapping regions. However, we identified two low frequency variants on chromosome 2 associated with CKD, one of them in the region of *SCG2* gene, which had been previously implicated with end-stage renal disease.^37^ These results illustrate the complementarity of the admixture and association approaches in identifying and improving the complex trait mapping in multi-ethnic populations.

There are some limitations of our findings. Our admixture mapping analysis captures both common and rare variants through local ancestry. However, the GWAS fine-mapping was limited to the genotype/imputed data and rare variants within the associated region were not interrogated. The available HL samples for validation had a different ancestry composition of the HCHS/SOL samples, which could explain in part the unsuccessfully validation in HLs.

In summary, we identified three novel loci associated with eGFR and CKD using admixture mapping in HL, which were not identified by the traditional GWAS approach. The signals on chromosomes 14 and 15 were driven by European and African-specific loci within the *DLK1-DIO3* imprinted domain and *RYR3* gene, respectively, and both replicated in WHI AA samples. The CKD signal on chromosome 2 was captured by an European-specific loci near the *FSHR* and *NRXN1* genes but the findings did not replicate. Our study provides new information into genetic contributions to eGFR in HL, and in particular, ancestry-specific genetic regions that are implicated with eGFR.

## Author Contributions

N. Franceschini and T. Thornton conceived and supervised the study. A. Horimoto analyzed the data and wrote the manuscript with input from all authors. J. Cai, J. Lash and M. Daviglus contributed to the final manuscript.

## Acknowledgments

The baseline examination of HCHS/SOL was carried out as a collaborative study supported by contracts from the National Heart, Lung, and Blood Institute (NHLBI) to the University of North Carolina (N01-HC65233), University of Miami (N01-HC65234), Albert Einstein College of Medicine (N01-HC65235), Northwestern University (N01-HC65236), and San Diego State University (N01-HC65237). The following Institutes/Centers/Offices contributed to the first phase of HCHS/SOL through a transfer of funds to the NHLBI: National Institute on Minority Health and Health Disparities, National Institute on Deafness and Other Communication Disorders, National Institute of Dental and Craniofacial Research (NIDCR), National Institute of Diabetes and Digestive and Kidney Diseases, National Institute of Neurological Disorders and Stroke, NIH Institution-Office of Dietary Supplements. The Genetic Analysis Center at Washington University was supported by NHLBI and NIDCR contracts (HHSN268201300005C AM03 and MOD03). Genotyping efforts were supported by NHLBI HSN 26220/20054C, NCATS CTSI grant UL1TR000124, and NIDDK Diabetes Research Center (DRC) grant DK063491. This study is supported by the following grants: National Institutes of Health DK117445, MD012765, HL140385 and HL123677 to N. Franceschini.

