## Supplemental for "Genome-wide admixture mapping of eGFR and CKD identify European and African ancestry-of-origin loci in U.S. Hispanics/Latinos"

**Table of Contents for the Supplemental Material**

**Supplemental Appendix 1**. Details on the admixture mapping linear and logistic mixed models used for eGFR and CKD analyses.

**Supplemental Figure 1**. Conditional admixture mapping analyses for eGFR and CKD including the lead SNP of each locus as covariate.

**Supplemental Figure 2.** Ancestry background of HCHS/SOL and WHI HL samples.

**Supplemental Table 1**. Annotation of the SNPs within ancestry-of-origin loci associated with eGFR and CKD.

**Supplemental Appendix 1. Details on the admixture mapping linear and logistic mixed models used for eGFR and CKD analyses.**

The admixture mapping for eGFR and CKD were performed using a joint test implemented in the GENESIS R package[1], in which European, African, and Native American ancestries were tested simultaneously. Overall, the analysis consisted of two steps. We first fit a mixed model under the null hypothesis of no genetic association, including random and fixed effects as detailed in the Methods section. The fitted null models are then used to test the association between the local ancestry at each locus and the outcomes. The local ancestry calls were estimated on the phased genotype data as described elsewhere[2]. For Hispanics/Latinos, it represents the European, African, and Native American allelic dosages (0, 1 or 2 copies of an ancestry-of-origin allele) at each locus.

For eGFR analysis, we used an admixture mapping linear mixed model. The full model is described by:

$$\boldsymbol{y} = \boldsymbol{X\alpha} + \boldsymbol{A_j\beta_j} + \boldsymbol{g} + \boldsymbol{\varepsilon},$$

where $\boldsymbol{y}$ is the vector of eGFR measures for the $N$ individuals, $\boldsymbol{X}$ is the vector of covariates, and $\boldsymbol{\alpha}$ is the vector of fixed covariate effects including an intercept. Letting the third ancestral population be the reference population, $\boldsymbol{A_j}$ represents a $Nx(K-1)$ matrix of the local ancestry allelic dosages at the locus $j$ for the $K-1$ parental populations, with the corresponding effect size vector $\boldsymbol{\beta_j}$ of length $K-1$. We assume that $\boldsymbol{g} \sim N(0, \sigma_a^2 \Phi)$ is a vector $\boldsymbol{g} = (g_1, \dots, g_N)$ of random effects for the $N$ subjects, where $\sigma_a^2$ is the additive genetic variance and $\Phi$ is the genetic relatedness matrix, and that $\varepsilon \sim N(0, \sigma_e^2 I)$ is a vector $\varepsilon = (\varepsilon_1, \dots, \varepsilon_N)$ of residual effects, where $\sigma_e^2$ represents the residual variance and $I$ is an identity matrix. The average Information Restricted Maximum Likelihood (AI-REML) approach was used to estimate the variance components $\sigma_a^2$ e $\sigma_e^2$ under the null model. The null hypothesis that $\boldsymbol{\beta_j} = \boldsymbol{0}$ was assessed via multivariate score test.

The admixture mapping linear mixed model was extended to analyze binary phenotypes, such as CKD, using an implementation that applies a penalized quasi-likelihood approximation to the generalized linear mixed model to fit the null model[3]. The full admixture logistic mixed model is described by:

$$logit(\pi) = \boldsymbol{X\alpha} + \boldsymbol{A_j\beta_j} + \boldsymbol{g},$$

where $\pi = P(y = 1|\boldsymbol{X}, \boldsymbol{A_j}, \boldsymbol{g})$ represents the $Nx1$ column vector of probabilities of being affected for the $N$ individuals conditional to covariates, local ancestry calls and random effects. $\boldsymbol{X}$, $\boldsymbol{\alpha}$, $\boldsymbol{A_j}$, $\boldsymbol{\beta_j}$ and $\boldsymbol{g}$ are defined as above. As for the eGFR analysis, we used a multivariate score test to assess the null hypothesis that $\boldsymbol{\beta_j} = \boldsymbol{0}$.

We also conducted secondary single ancestry admixture mapping analyses to identify which ancestry population was driving the signal in each associated locus. Here, we tested each ancestry against the others to assess the effect of each ancestry separately. The local ancestry allelic dosages of the reference population (European, for example) is included as a predictor in the model and compared to the non-reference group (African + Native American in the example).

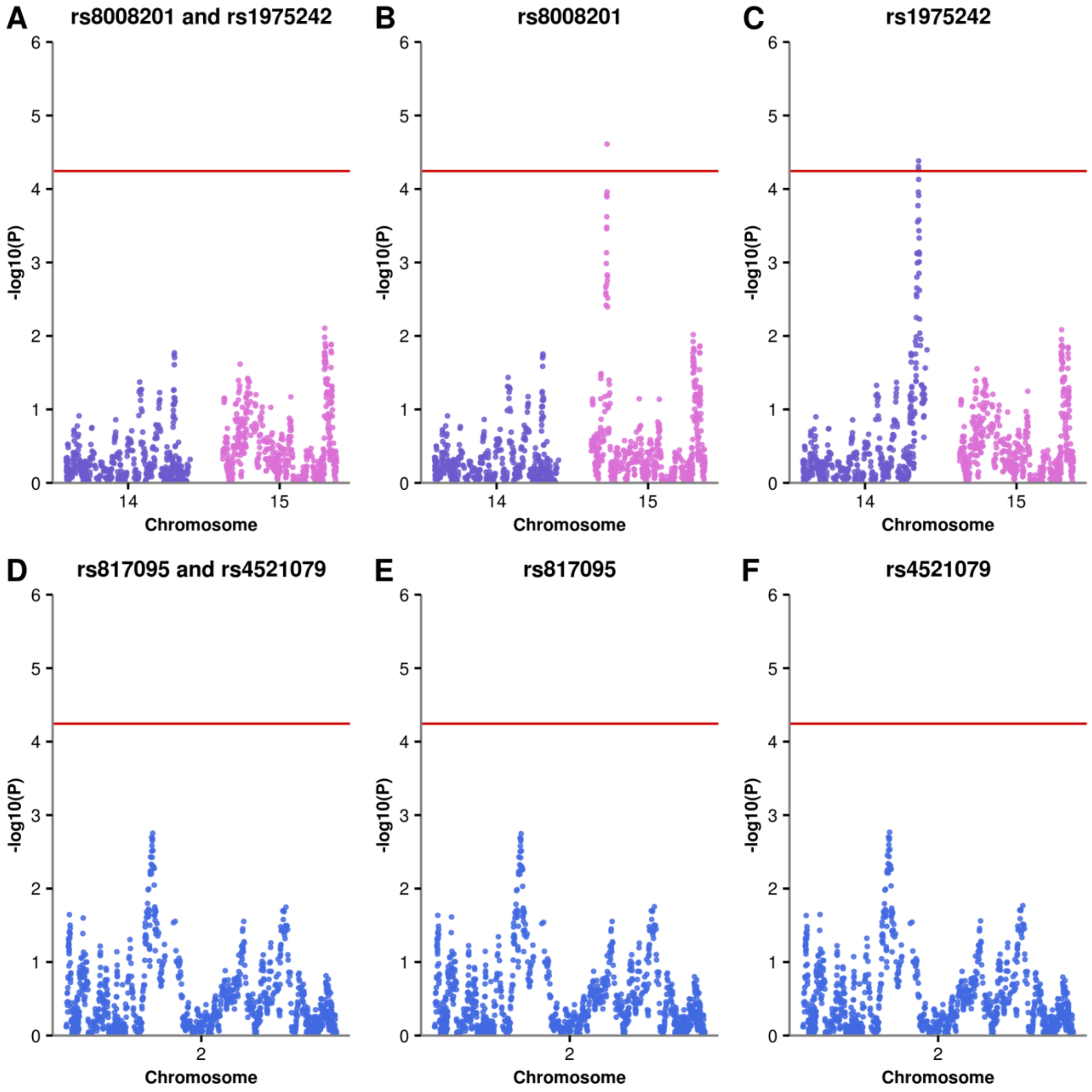


**Supplemental Figure 1**. **Conditional admixture mapping analyses for eGFR and CKD including the lead SNP of each locus as covariate.** eGFR: (**A**) both SNPs from chromosomes 14 (rs8008201) and 15 (rs1975242) loci; (**B**) SNP from chromosome 14 locus (rs8008201); (**C**) SNP from chromosome 15 locus (rs1975242). CKD: (**D**) SNPs from both loci on chromosome 2 (rs817095 and rs4521079); (**E**) SNP rs817095 from the first locus; (**F**) SNP rs4521079 from the second locus. Note that adjusting for SNPs within chromosome 14 does not reduce the signal for chromosome 15, and vice-versa as seen in **B** and **C**.

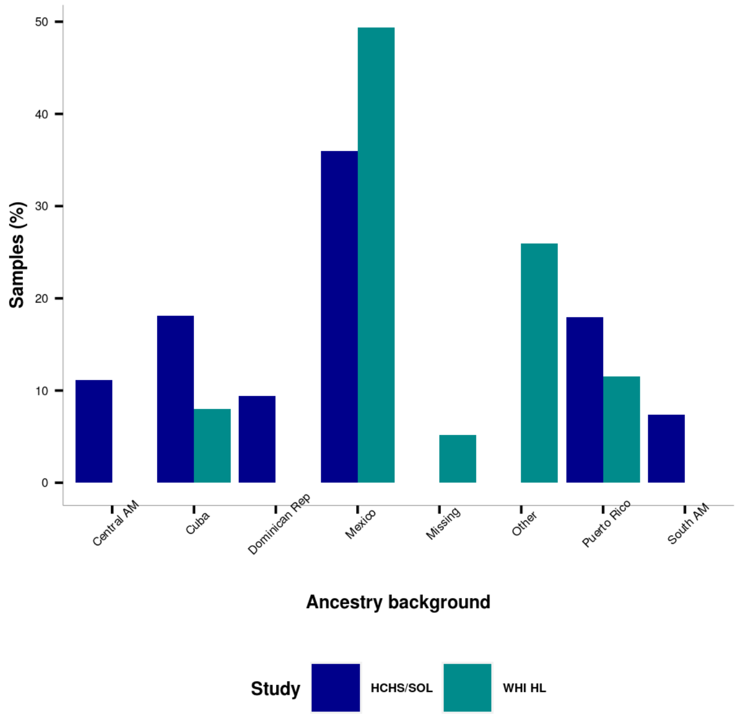


**Supplemental Figure 2**. **Ancestry background of HCHS/SOL and WHI HL samples.** Central AM: Central America; Dominican Rep: Dominican Republic; South AM: South America.

**Supplemental Table 1**. **Annotation of the SNPs within ancestry-of-origin loci associated with eGFR and CKD.**

| SNP | Chr | Phypos | Ref | Alt | AFR.f | EUR.f | NAM.f | Gene / Symbol | Feature type | Consequence |
|---|---|---|---|---|---|---|---|---|---|---|
| rs817095 | 2 | 49571948 | G | A | 0.93 | 0.80 | 0.85 | --- | --- | intergenic |
| rs860133 | 2 | 49578937 | T | G | 0.50 | 0.18 | 0.33 | --- | --- | intergenic |
| rs860994 | 2 | 49580566 | G | A | 0.05 | 0.09 | 0.04 | --- | --- | intergenic |
| rs817081 | 2 | 49588657 | T | G | 0.37 | 0.12 | 0.12 | --- | --- | intergenic |
| rs7583216 | 2 | 49599540 | G | A | 0.27 | 0.09 | 0.23 | --- | --- | intergenic |
| rs2882306 | 2 | 49620211 | C | T | 0.64 | 0.80 | 0.88 | --- | --- | intergenic |
| rs817035 | 2 | 49623006 | C | T | 0.29 | 0.17 | 0.27 | --- | --- | intergenic |
| rs7580237 | 2 | 49623299 | G | T | 0.04 | 0.00 | 0.00 | --- | --- | intergenic |
| rs817038 | 2 | 49626781 | G | A | 0.80 | 0.90 | 0.93 | --- | --- | intergenic |
| rs17038697 | 2 | 49628461 | G | A | 0.06 | 0.00 | 0.04 | --- | --- | intergenic |
| rs1882345 | 2 | 49630281 | T | C | 0.56 | 0.73 | 0.66 | --- | --- | intergenic |
| rs817043 | 2 | 49630633 | T | G | 0.13 | 0.10 | 0.06 | --- | regulatory | regulatory |
| rs698840 | 2 | 49635681 | A | G | 0.66 | 0.83 | 0.72 | --- | --- | intergenic |
| rs817060 | 2 | 49639121 | A | G | 0.04 | 0.09 | 0.04 | --- | --- | intergenic |
| rs10170288 | 2 | 49640979 | G | A | 0.39 | 0.64 | 0.39 | --- | --- | intergenic |
| rs1527898 | 2 | 49641262 | T | G | 0.31 | 0.19 | 0.19 | --- | --- | intergenic |
| rs11125238 | 2 | 49643063 | C | A | 0.35 | 0.28 | 0.37 | --- | --- | intergenic |
| rs11890365 | 2 | 49643222 | G | A | 0.06 | 0.00 | 0.01 | --- | --- | intergenic |
| rs17038751 | 2 | 49644800 | A | G | 0.24 | 0.11 | 0.13 | --- | --- | intergenic |
| rs11892442 | 2 | 49646196 | T | C | 0.12 | 0.00 | 0.01 | --- | --- | intergenic |
| rs10172488 | 2 | 49646785 | G | A | 0.21 | 0.22 | 0.13 | --- | --- | intergenic |
| rs1405959 | 2 | 49647212 | A | G | 0.74 | 0.50 | 0.57 | --- | --- | intergenic |
| rs843840 | 2 | 49650145 | A | G | 0.23 | 0.11 | 0.07 | --- | --- | intergenic |
| rs12105450 | 2 | 49653672 | C | T | 0.46 | 0.33 | 0.27 | --- | --- | intergenic |
| rs1405969 | 2 | 49676328 | A | G | 0.55 | 0.61 | 0.57 | --- | --- | intergenic |
| rs10495974 | 2 | 49678339 | C | T | 0.31 | 0.24 | 0.40 | --- | --- | intergenic |
| rs1405955 | 2 | 49684567 | T | C | 0.90 | 0.97 | 0.97 | --- | --- | intergenic |
| rs10186177 | 2 | 49703199 | A | C | 0.21 | 0.46 | 0.37 | --- | --- | intergenic |
| rs13427322 | 2 | 49707932 | G | A | 0.07 | 0.07 | 0.03 | --- | --- | intergenic |

| | | | | | | | | | | |
|---|---|---|---|---|---|---|---|---|---|---|
| rs11888995 | 2 | 49714129 | T | C | 0.70 | 0.73 | 0.76 | --- | --- | intergenic |
| rs13382313 | 2 | 49714846 | C | T | 0.12 | 0.37 | 0.32 | --- | --- | intergenic |
| rs12987465 | 2 | 49715021 | G | A | 0.58 | 0.36 | 0.44 | --- | --- | intergenic |
| rs977134 | 2 | 49716202 | C | T | 0.21 | 0.08 | 0.22 | --- | --- | intergenic |
| rs1405966 | 2 | 49716853 | A | G | 0.78 | 0.79 | 0.80 | --- | --- | intergenic |
| rs12996690 | 2 | 49724966 | T | C | 0.19 | 0.28 | 0.21 | --- | --- | intergenic |
| rs1405965 | 2 | 49737819 | A | G | 0.79 | 0.73 | 0.61 | --- | --- | intergenic |
| rs6545119 | 2 | 49739361 | C | T | 0.77 | 0.83 | 0.72 | --- | --- | intergenic |
| rs13422448 | 2 | 49743324 | A | C | 0.01 | 0.32 | 0.24 | --- | --- | intergenic |
| rs6709183 | 2 | 49749038 | C | T | 0.50 | 0.24 | 0.40 | --- | --- | intergenic |
| rs4971621 | 2 | 49752589 | T | C | 0.15 | 0.53 | 0.48 | --- | --- | intergenic |
| rs17038952 | 2 | 49788135 | A | C | 0.09 | 0.05 | 0.03 | --- | --- | intergenic |
| rs1995172 | 2 | 49790063 | C | T | 0.19 | 0.57 | 0.46 | --- | --- | intergenic |
| rs17038971 | 2 | 49795342 | G | A | 0.06 | 0.00 | 0.01 | --- | --- | intergenic |
| rs7595953 | 2 | 49796689 | A | G | 0.19 | 0.05 | 0.04 | --- | --- | intergenic |
| rs13429217 | 2 | 49808562 | C | T | 0.21 | 0.09 | 0.21 | --- | --- | intergenic |
| rs12469706 | 2 | 49811212 | T | C | 0.01 | 0.07 | 0.03 | --- | --- | intergenic |
| rs1593705 | 2 | 49816534 | G | A | 0.29 | 0.73 | 0.61 | --- | --- | intergenic |
| rs13399003 | 2 | 49819514 | G | T | 0.20 | 0.05 | 0.04 | --- | --- | intergenic |
| rs13401858 | 2 | 49825768 | G | A | 0.01 | 0.24 | 0.22 | --- | regulatory | regulatory |
| rs1553129 | 2 | 49825824 | C | T | 0.09 | 0.00 | 0.01 | --- | --- | intergenic |
| rs1498799 | 2 | 49829630 | A | G | 0.03 | 0.34 | 0.24 | --- | --- | intergenic |
| rs2162518 | 2 | 49835794 | A | C | 0.25 | 0.11 | 0.23 | --- | --- | intergenic |
| rs17039077 | 2 | 49847520 | G | A | 0.05 | 0.00 | 0.01 | --- | --- | intergenic |
| rs13423597 | 2 | 49849103 | A | G | 0.11 | 0.32 | 0.36 | --- | --- | intergenic |
| rs7577053 | 2 | 49849708 | T | C | 0.62 | 0.74 | 0.74 | --- | --- | intergenic |
| rs1391748 | 2 | 49850450 | C | T | 0.15 | 0.32 | 0.37 | --- | --- | intergenic |
| rs4527244 | 2 | 49855331 | T | C | 0.92 | 0.83 | 0.86 | --- | --- | intergenic |
| rs976704 | 2 | 49856326 | C | T | 0.76 | 0.50 | 0.50 | --- | --- | intergenic |
| rs13425236 | 2 | 49858261 | T | C | 0.08 | 0.31 | 0.35 | --- | --- | intergenic |
| rs11125265 | 2 | 49861898 | T | C | 0.82 | 0.62 | 0.54 | --- | --- | intergenic |
| rs7564547 | 2 | 49862134 | A | C | 0.92 | 0.90 | 0.89 | --- | --- | intergenic |
| rs2350699 | 2 | 49871330 | G | A | 0.07 | 0.10 | 0.09 | --- | --- | intergenic |
| rs17794707 | 2 | 49876591 | A | C | 0.03 | 0.25 | 0.16 | --- | --- | intergenic |

| | | | | | | | | | | |
|---|---|---|---|---|---|---|---|---|---|---|
| rs12713068 | 2 | 49887106 | A | G | 0.17 | 0.40 | 0.44 | --- | --- | intergenic |
| rs6545125 | 2 | 49888676 | G | A | 0.74 | 0.51 | 0.47 | --- | --- | intergenic |
| rs10190188 | 2 | 49894154 | T | C | 0.11 | 0.34 | 0.24 | --- | --- | intergenic |
| rs4521079 | 2 | 49909155 | G | A | 0.79 | 0.58 | 0.73 | --- | --- | intergenic |
| rs1568287 | 2 | 49909547 | C | A | 0.75 | 0.18 | 0.31 | --- | --- | intergenic |
| rs2176603 | 2 | 49909670 | A | C | 0.23 | 0.07 | 0.08 | --- | --- | intergenic |
| rs2350701 | 2 | 49913955 | G | A | 0.58 | 0.46 | 0.52 | --- | --- | intergenic |
| rs10185615 | 2 | 49944565 | A | C | 0.08 | 0.40 | 0.26 | --- | --- | intergenic |
| rs7593733 | 2 | 49963077 | C | A | 0.42 | 0.75 | 0.50 | --- | --- | intergenic |
| rs17489439 | 2 | 49964349 | G | A | 0.04 | 0.34 | 0.21 | --- | --- | intergenic |
| rs13386956 | 2 | 49964349 | T | C | 0.21 | 0.00 | 0.02 | --- | --- | intergenic |
| rs2139156 | 2 | 49972394 | T | C | 0.04 | 0.44 | 0.27 | --- | --- | intergenic |
| rs1914782 | 2 | 49975135 | C | T | 0.11 | 0.34 | 0.23 | --- | --- | intergenic |
| rs10171772 | 2 | 49977093 | G | A | 0.37 | 0.44 | 0.45 | --- | --- | intergenic |
| rs1914779 | 2 | 49986278 | C | T | 0.43 | 0.08 | 0.27 | --- | --- | intergenic |
| rs7349353 | 2 | 49990017 | A | G | 0.12 | 0.32 | 0.22 | --- | --- | intergenic |
| rs1518823 | 2 | 49990368 | C | T | 0.96 | 0.89 | 0.81 | --- | --- | intergenic |
| rs7595014 | 2 | 49992053 | A | G | 0.62 | 0.36 | 0.47 | --- | --- | intergenic |
| rs870168 | 2 | 49997218 | G | A | 0.88 | 0.45 | 0.66 | --- | --- | intergenic |
| rs11903512 | 2 | 50005959 | T | C | 0.19 | 0.03 | 0.05 | --- | --- | intergenic |
| rs17039294 | 2 | 50009220 | A | G | 0.04 | 0.01 | 0.01 | --- | --- | intergenic |
| rs17039309 | 2 | 50011821 | G | A | 0.14 | 0.00 | 0.01 | --- | --- | intergenic |
| rs17039328 | 2 | 50022031 | T | C | 0.51 | 0.31 | 0.26 | --- | --- | intergenic |
| rs7586200 | 2 | 50031813 | G | A | 0.32 | 0.02 | 0.04 | --- | --- | intergenic |
| rs1518834 | 2 | 50048977 | C | A | 0.16 | 0.15 | 0.10 | --- | --- | intergenic |
| rs885560 | 2 | 50055938 | A | G | 0.02 | 0.16 | 0.09 | --- | --- | intergenic |
| rs925931 | 2 | 50059808 | T | C | 0.07 | 0.16 | 0.10 | --- | --- | intergenic |
| rs12466419 | 2 | 50061043 | A | G | 0.03 | 0.22 | 0.28 | --- | --- | intergenic |
| rs1363047 | 2 | 50085965 | A | G | 0.08 | 0.19 | 0.12 | --- | --- | intergenic |
| rs1156742 | 2 | 50125840 | G | A | 0.62 | 0.51 | 0.56 | --- | --- | intergenic |
| rs1001943 | 2 | 50127552 | T | C | 0.23 | 0.01 | 0.04 | --- | --- | intergenic |
| rs7558063 | 2 | 50131767 | A | C | 0.46 | 0.12 | 0.14 | --- | --- | intergenic |
| rs971732 | 2 | 50142117 | A | C | 0.41 | 0.37 | 0.42 | NRXN1 | transcript | downstream |
| rs1045881 | 2 | 50148972 | C | T | 0.12 | 0.16 | 0.10 | NRXN1 | transcript | 3 prime UTR |

| | | | | | | | | | | |
|---|---|---|---|---|---|---|---|---|---|---|
| | | | | | | | | NRXN1<br>NRXN1 | transcript<br>transcript | downstream<br>non-coding exon |
| rs17039448 | 2 | 50149510 | A | G | 0.22 | 0.19 | 0.30 | NRXN1<br>NRXN1 | transcript<br>transcript | intron<br>non-coding |
| rs1421594 | 2 | 50176965 | G | A | 0.48 | 0.38 | 0.43 | NRXN1<br>NRXN1 | transcript<br>transcript<br>regulatory | intron<br>non-coding<br>regulatory |
| rs2193870 | 2 | 50179975 | C | T | 0.01 | 0.24 | 0.20 | NRXN1<br>NRXN1 | transcript<br>transcript | intron<br>non-coding |
| rs17491881 | 2 | 50188135 | T | C | 0.10 | 0.19 | 0.12 | NRXN1<br>NRXN1 | transcript<br>transcript | intron<br>non-coding |
| rs17439140 | 2 | 50195048 | C | T | 0.13 | 0.34 | 0.38 | NRXN1<br>NRXN1 | transcript<br>transcript | intron<br>non-coding |
| rs11125280 | 2 | 50197075 | C | T | 0.52 | 0.28 | 0.40 | NRXN1<br>NRXN1 | transcript<br>transcript | intron<br>non-coding |
| rs8008201 | 14 | 101329926 | G | A | 0.09 | 0.23 | 0.12 | MEG3<br>RP11-123M6.2 | transcript<br>transcript | downstream<br>upstream |
| rs8021312 | 14 | 101333646 | T | C | 0.32 | 0.25 | 0.20 | MIR493 | transcript | upstream |
| rs11851174 | 14 | 101345504 | C | T | 0.37 | 0.21 | 0.16 | MIR337<br>RTL1<br>MIR665<br>MIR433<br>MIR127<br>MIR431 | transcript<br>transcript<br>transcript<br>transcript<br>transcript<br>transcript | downstream<br>downstream<br>downstream<br>upstream<br>upstream<br>upstream |
| rs3825569 | 14 | 101350298 | T | C | 0.61 | 0.64 | 0.43 | MIR433<br>MIR127<br>MIR431<br>MIR136<br>MIR432<br>RTL1 | transcript<br>transcript<br>transcript<br>transcript<br>transcript<br>transcript | downstream<br>downstream<br>downstream<br>upstream<br>upstream<br>synonymous |
| rs1975242 | 15 | 33801207 | G | A | 0.67 | 0.49 | 0.32 | RYR3 | transcript | intron |
| rs7165389 | 15 | 33801946 | T | C | 0.21 | 0.13 | 0.09 | RYR3 | transcript | intron |

| | | | | | | | | | | |
|---|---|---|---|---|---|---|---|---|---|---|
| rs2596211 | 15 | 33804910 | T | G | 0.98 | 0.66 | 0.55 | RYR3 | transcript | intron |
| rs11632989 | 15 | 33806622 | C | T | 0.00 | 0.11 | 0.07 | RYR3 | transcript | intron |
| rs6495130 | 15 | 33810168 | G | A | 0.44 | 0.33 | 0.20 | RYR3 | transcript | intron |
| rs735545 | 15 | 33811083 | T | G | 0.57 | 0.46 | 0.29 | RYR3 | transcript regulatory | intron regulatory |
| rs4281677 | 15 | 33816995 | C | T | 0.47 | 0.12 | 0.12 | RYR3 | transcript | intron |
| rs2596220 | 15 | 33818190 | G | A | 0.41 | 0.36 | 0.52 | RYR3 | transcript | intron |
| rs2572203 | 15 | 33818341 | C | T | 0.92 | 0.85 | 0.86 | RYR3 | transcript | intron |
| rs12909478 | 15 | 33829250 | C | T | 0.09 | 0.20 | 0.11 | RYR3 | transcript | intron |
| rs957467 | 15 | 33830326 | T | C | 0.44 | 0.71 | 0.73 | RYR3 | transcript | intron |
| rs6495164 | 15 | 33838470 | C | T | 0.11 | 0.17 | 0.33 | RYR3 | transcript | intron |
| rs1435102 | 15 | 33839194 | A | G | 0.46 | 0.82 | 0.80 | RYR3 | transcript | intron |
| rs7163719 | 15 | 33841956 | T | C | 0.28 | 0.01 | 0.02 | RYR3 | transcript | intron |
| rs1865495 | 15 | 33847805 | T | C | 0.26 | 0.53 | 0.62 | RYR3 | transcript | intron |
| rs9744361 | 15 | 33857949 | C | A | 0.55 | 0.12 | 0.12 | RYR3 | transcript | intron |
| rs1435118 | 15 | 33865455 | G | A | 0.34 | 0.74 | 0.73 | RYR3 | transcript | intron |
| rs8041171 | 15 | 33866510 | C | A | 0.14 | 0.03 | 0.04 | RYR3 | transcript | intron |
| rs680851 | 15 | 33866632 | T | C | 0.34 | 0.74 | 0.73 | RYR3 | transcript | intron |
| rs674155 | 15 | 33872177 | C | T | 0.40 | 0.77 | 0.76 | RYR3 | transcript | missense splice synonymous |
| rs683484 | 15 | 33872692 | G | A | 0.40 | 0.77 | 0.76 | RYR3 | transcript regulatory | intron regulatory |
| rs581954 | 15 | 33873369 | A | C | 0.13 | 0.14 | 0.12 | RYR3 | transcript regulatory | intron regulatory |
| rs668570 | 15 | 33873657 | C | T | 0.87 | 0.81 | 0.61 | RYR3 | transcript regulatory | intron regulatory |
| rs2077268 | 15 | 33873751 | G | A | 0.36 | 0.11 | 0.07 | RYR3 | transcript regulatory | missense variant regulatory |
| rs748298 | 15 | 33873901 | C | T | 0.41 | 0.07 | 0.10 | RYR3 | transcript regulatory | intron regulatory |
| rs3794586 | 15 | 33875264 | G | T | 0.07 | 0.12 | 0.24 | RYR3 | transcript | intron |
| rs10431811 | 15 | 33876449 | G | A | 0.34 | 0.46 | 0.44 | RYR3 | transcript | intron |
| rs2643364 | 15 | 33876517 | A | G | 0.88 | 0.75 | 0.65 | RYR3 | transcript | intron |

| | | | | | | | | | | |
|---|---|---|---|---|---|---|---|---|---|---|
| rs659517 | 15 | 33877066 | G | T | 0.31 | 0.10 | 0.25 | RYR3 | transcript<br>regulatory | intron<br>regulatory |
| rs2643363 | 15 | 33877938 | A | G | 0.21 | 0.23 | 0.13 | RYR3 | transcript | intron |
| rs16972835 | 15 | 33878846 | C | A | 0.12 | 0.10 | 0.08 | RYR3 | transcript | intron |
| rs16972837 | 15 | 33879387 | G | A | 0.19 | 0.10 | 0.21 | RYR3 | transcript | intron |
| rs10153042 | 15 | 33879608 | A | G | 0.53 | 0.38 | 0.28 | RYR3 | transcript | intron |
| rs671122 | 15 | 33879678 | C | T | 0.84 | 0.64 | 0.57 | RYR3 | transcript | intron |
| rs658750 | 15 | 33880181 | G | A | 0.65 | 0.54 | 0.36 | RYR3 | transcript | intron |
| rs688939 | 15 | 33880493 | A | G | 0.04 | 0.13 | 0.06 | RYR3 | transcript | intron |
| rs640152 | 15 | 33884488 | A | G | 0.84 | 0.64 | 0.58 | RYR3 | transcript<br>regulatory | intron<br>regulatory |

Chr: chromosome; Phypos: physical position in build GRCh37/hg19; Ref: reference allele (forward strand); Alt: alternative allele (forward strand); AFR.f: alternative allele frequency in African populations; EUR.f: alternative allele frequency in European populations; NAM.f: alternative allele frequency in Native American populations. Both reference allele and population-specific reference allele frequency were obtained from 1000 Genomes Project phase 3 using Ensembl (https://uswest.ensembl.org/index.html); Gene/Symbol, Feature type and Consequence were obtained using the Ensembl Variant Effect Predictor tool (https://uswest.ensembl.org/info/docs/tools/vep/index.html).
